# From Sequence to Function: Coevolving Amino Acids Encode Structural and Functional Domains

**DOI:** 10.1101/109397

**Authors:** Daniele Granata, Luca Ponzoni, Cristian Micheletti, Vincenzo Carnevale

## Abstract

Amino acids interactions within protein families are so optimized that the sole analysis of evolutionary co-mutations can identify pairs of contacting residues. It is also known that evolution conserves functional dynamics, i.e., the concerted motion or displacement of large protein regions or domains. Is it, therefore, possible to use a pure sequence-based analysis to identify these dynamical domains? To address this question, we introduce here a general co-evolutionary coupling analysis strategy and apply it to a curated sequence database of hundreds of protein families. For most families, the sequence-based method partitions amino acids into few clusters. When viewed in the context of the native structure, these clusters have the signature characteristics of viable protein domains: they are spatially separated but individually compact. They have a direct functional bearings too, as shown for various reference cases. We conclude that even large-scale structural and functionally-related properties can be recovered from inference methods applied to evolutionary-related sequences. The method introduced here is available as a software package and web server (http://spectrus.sissa.it/spectrus-evo_webserver).

## Introduction

A powerful paradigm and organizing principle in molecular biology is the flow of information from the chemical composition to the biological functionality of proteins. The flow is typically viewed as a chain of implications: the protein sequence encodes for the structure, which, in turn, underpins function.

Accordingly, much attention has been, and still is, paid to the two relationships in the tripartite ladder: *sequence-structure* and *structure-function.* These, however, have mostly been treated as distinct problems, each to be addressed within its own conceptual framework, and with *ad hoc* tools.

For instance, an increasingly credited mediator of structure and biological function is the internal dynamics of proteins. In fact, the secondary and higher-order organization of several enzymes endows them with innate capabilities to sustain large-scale conformational changes that are functionally oriented and can be excited by thermal fluctuations or triggered by the binding of ligands and effectors [1–17]. Our current understanding of these mechanisms has been much shaped by single-molecule experiments that have provided vivid and quantitative descriptions of the dynamical basis of protein function [18–22]. Computational and theoretical approaches, from atomistic molecular dynamics (MD) simulations [23–25] to coarse-grained elastic networks [26–28], have also provided a detailed understanding of how the structural architecture reverberates in the internal dynamics. These approaches have clarified that the structural changes that accompany and assist function can be well captured by only few collective degrees of freedom, including those resulting from the relative movement of quasi-rigid (dynamical) domains [29–32].

Likewise, the theoretical exploration of the sequence-structure relationship [33, 34] has also been devoted to devise methods for predicting the structure of a protein from sequence alone [35–37], with recent methodological breakthroughs due to the application of statistical inference techniques to the analysis of multiple sequence alignments. Correlated substitutions can help identify those sites that host co-evolving mutations and these, in turn, are an indicator of spatial proximity [38–46].

The natural question posed by these parallel, separate advancements in the sequence-structure and structure-function realms is whether or not it is at all feasible to establish a more direct connection between them [47]. In particular, one may ask whether coevolutionary couplings inferred from multiple sequence analysis can be related to functional properties in a seamless manner, that is without relying on the prior knowledge of the structure.

To our knowledge, this overarching question has been addressed in few studies tackling it in specific contexts [48–58], but not yet in a general manner.

Here, motivated by these studies and especially by the protein sector analysis of ref. [49], we carry out a first systematic characterization of the sequence → function relationship, by using sequence-based coevolutionary data to infer dynamical/functional domains whose organization has been conserved along evolution. We term such fundamental units “evolutionary domains”.

Specifically, we first survey the coevolution-based profiling of functional domains for a previously-annotated set of about 800 multiple sequence alignments [43]. From this extensive survey we find that domains inferred from the sole sequence-based, coevolutionary analysis are compact in space and well consistent with the dynamical, or quasi-rigid domains inferred from the analysis of small and large-scale structural fluctuations. After this general, database-wide application, we next move to a more in-depth discussion of a biophysically-relevant case of three distinct subfamilies of the ion channels superfamily [59]. We show how the same structural template, inherited from a common ancestor gene, can give rise to distinct sets of evolutionary domains, thus recapitulating the major functional differences observed for the three subfamilies, demonstrating the relevance of the method for comparative analysis even in presence of few sequences.

## Results and Discussion

Our goal is to use extensive sets of mutual statistical dependencies in the patterns of mutations within a protein family, to identify groups of residues that have arguably evolved concertedly because of structural and functional constraints. We refer to these groups or clusters of amino acids as evolutionary domains (EDs). To devise an “evolutionary informed” criterion for such clustering purpose, we introduce a similarity or proximity measure between residues based on the pairwise correlations identified in multiple sequence alignments (MSA). The subdivision of a sequence in multiple EDs is then the result of a clustering procedure, the spectral clustering, which returns an optimal set of densely connected groups.

## From correlated mutations to evolutionary domains

We adopt a clustering strategy similar to that used in SPECTRUS [32], an algorithm to efficiently determine dynamical, quasi-rigid domains in protein complexes. In that case, the similarity measure was derived from reside-residue distance fluctuations and the clustering produced a hierarchy of subdivisions grouping together spatially compact residues sharing a common overall motion pattern.

Here we start from an MSA and the corresponding matrix of statistical couplings, **J**, calculated with one of the available coevolutionary analysis methods. Our method of choice for this step is the plmDCA approach described in ref. [43]; we discuss other methods, such as gplmDCA and plmDCA20, in SI to show that the definition of EDs is robust and does not depend on the approximation chosen to perform the statistical inference. We interpret each statistical coupling *J_ij_*as a measure of evolutionary proximity between residues *i* and *j* and use this identification to generate a similarity matrix **S**, to be used in a clustering algorithm (see workflow in Fig. 1a). In particular, these similarities are used to generate a sparse, undirected graph *G*(*V,E*), in which the vertices *V* correspond to the residues and the (weighted) edges *E* are given by the matrix elements *S_ij_*. Additionally, among the edges connected with each vertex, we retain only the top *k* = 7 strongest couplings, and then symmetrize the graph. The neighbour-restriction criterion ensures the robustness of the subsequent clustering, which is performed with a spectral clustering scheme [60], see Methods for more details. Finally a quality score, function of the number of clusters Q, is used to identify the most meaningful subdivisions, which are then taken to define the EDs.

**Figure 1.**
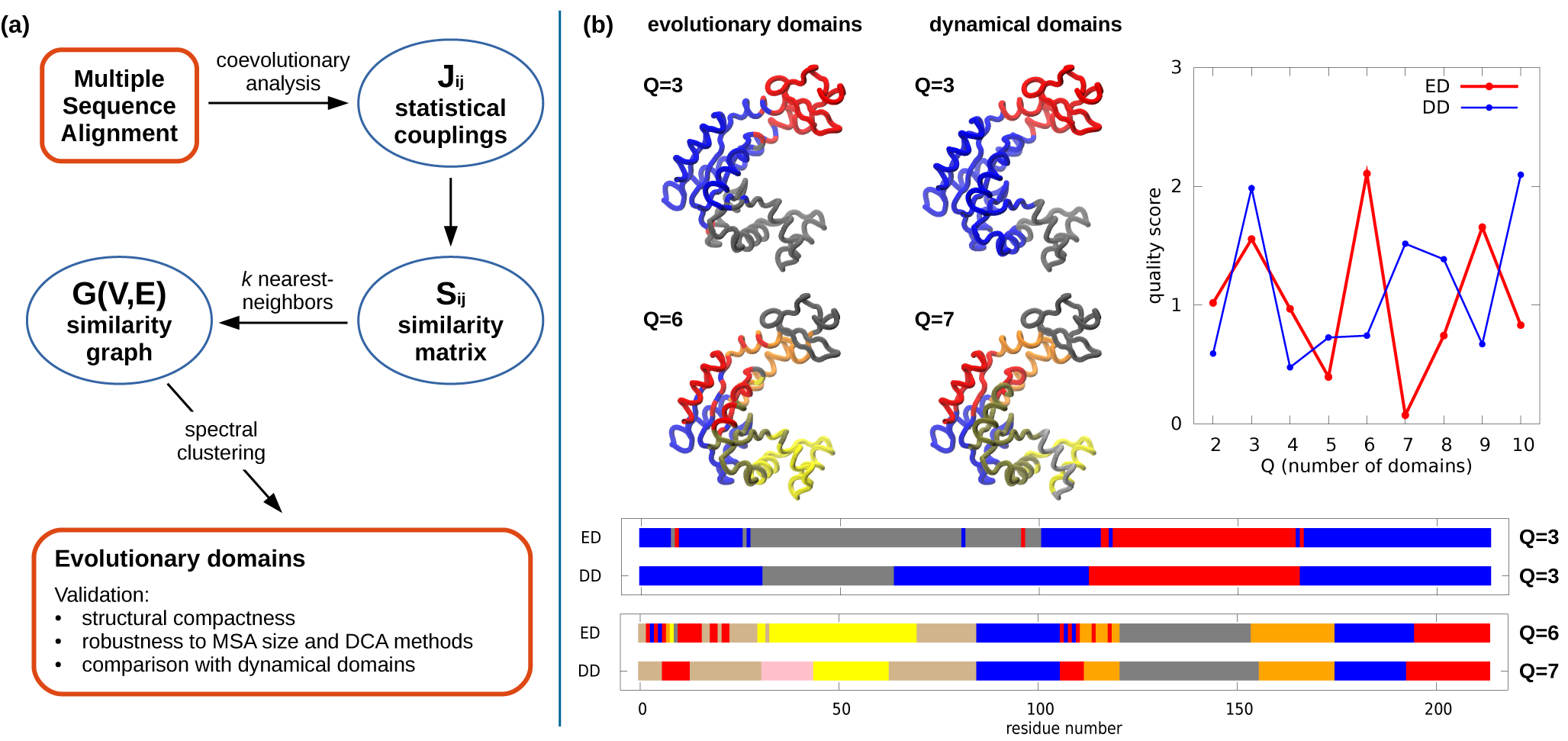
Left: schematic illustration of the steps performed for identifying the groups of coevolving residues (EDs: evolutionary domains) from a protein multiple sequence alignment. Right: analysis of the emblematic case of adenylate kinase. The evolutionary partitioning is contrasted with the subdivisions into quasi-rigid, dynamical domains obtained from the analysis of an MD simulation with the SPECTRUS webserver [32]. For the main sequence-and dynamics-based subdivisions, characterized by a high quality score, two color-coded representations, on the protein structure and on its sequence, are provided.

## A test case: adenylate kinase

To illustrate and validate the sequence-based ED decomposition, we first apply it to *Escherichia coli* adenylate kinase (PDBID: 4AKE), a standard benchmark for domain partitioning methods. The results are given in Fig. 1b. The quality score profile indicates that the best sequence-based partitionings are found for *Q* = 3, 6 and 9 domains, some of which are shown in Fig. 1b. We note that the EDs correspond to regions that are compact from the structural point of view, in a non-trivial way, involving specific secondary motifs and tertiary contacts.

This is a notable and intriguing result because the subdivisions are exclusively based on sequence, with no input about spatial organization. Therefore this compactness suggests that EDs might highlight those structural modules whose integrity was preserved along evolution. The division into 3 clusters is in this respects instructive as it appears consistent with the functional role of the two lateral flaps of the kinase, which are the binding sites for ATP (domain in red) and AMP (in gray) for the reversible catalysis of ADP.

To further clarify this point, we compared the EDs to the dynamical domains (DDs) resulting from a decomposition of adenylate kinase into quasi-rigid domains [32]. For this purpose we analyzed the conformers explored during two long molecular dynamics simulations started from either the open or the closed state. The DDs, which are inherently structure-based, are shown in Fig. 1b. Despite the very diverse nature of the input data, the correspondence between EDs and DDs is apparent at all levels of spatial resolution. Indeed, even the finest partitioning (*Q* = 9, Fig. S1) provides consistent decompositions in the two cases and highlights structural elements that are arguably crucial for the protein dynamics.

## Database-wide survey

For a more systematic characterization of the EDs, we extended the analysis to a dataset of 813 multiple sequence alignments compiled by Feinauer *et al.*[43]. This set is largely based on Pfam families [61] and was chosen for several reasons. First, it gives a comprehensive coverage of protein lengths, from 30 to 500 amino acids andnumber of sequences contained in the MSAs, from 16 to 65,000. Second, a reference PDB structure is available for each of the MSAs. The availability of these structural representatives, while not needed as input for the ED analysis (which is sequence-based only), is required for assessing *a posteriori* the spatial compactness of the EDs and their relationship with quasi-rigid or dynamical domains.

### Clustering Propensity and Community Structure of the Network of Couplings

As a preliminary study, we first investigated if the input network of statistical couplings *J_ij_*, obtained from coevolutionary analysis, exhibits an intrinsic propensity to be densely organized and, thus, to be clustered. Starting from a similarity graph *G_k_*(*V,E*) of each MSA, in which only the *k* strongest couplings are retained, it is possible to evaluate this propensity by calculating the weighted clustering coefficient *C* of the graph [62] and using a corresponding randomized graph as a reference (∆*C*=*C*–*C_rand_*). This quantity measures the probability that two neighbors of a vertex are also connected between themselves. As described in detail in the Methods section, this measure proves useful also to choose the optimal values of *k*, since, especially for MSA containing a large number of sequences, the different graphs show usually a maximum in the clustering coefficient at *k* = 7. Importantly, the MSA size (calculated as effective number of sequences, 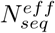, *i.e.* the number of sequences in the set whose maximum mutual identity is 90%) crucially affects the clustering propensity of the similarity graph, as clarified by the strong correlation between these two quantities shown in fig. S4. Thus, when a large dataset is available and the reconstruction of the network of couplings is most reliable, the latter shows a high tendency to cluster and an unambiguous number of “relevant” neighbors (*k*= 7), which is indicative of an inherent collective organization of the coevolution patterns. This differs significantly from the contact prediction problem, for which the number of neighbors to keep in order to maximize the true discovery rate is, on average, one per residue [41–43]. Strikingly, this number coincides with the average structural neighbors surrounding each residue in protein structures (6.75 ±0.04, calculated on the PDB structures of this dataset using a *C*_β_-*C*_β_ distance threshold of 8.5 Åas in [43]).

### Compactness of Evolutionary Domains

We then evaluated the spatial compactness of EDs for the 813 MSAs, to ascertain if the communities identified in the network of couplings are also structurally localized. This property is summarized by the quantity Ω, which is based on the number of residues that are spatially isolated, *i.e.* with a C_α_-C_α_ distance larger than10 AÅ from any other residue of the cluster (see Methods for details). We thus analyzed the entire dataset using two distinct summary statistics: i) the compactness of each MSA averaged over all the possible partitionings, *i.e.* from *Q* = 2 to *Q* = 10, 〈Ω*_α_*〉*_Q_* (Fig. 2, S5 and S6); ii) the average compactness of a partition characterized by *Q* clusters (averaged over the distinct MSAs), 〈Ω^Q^〉_*α*_ (Fig. S7).

**Figure 2.**
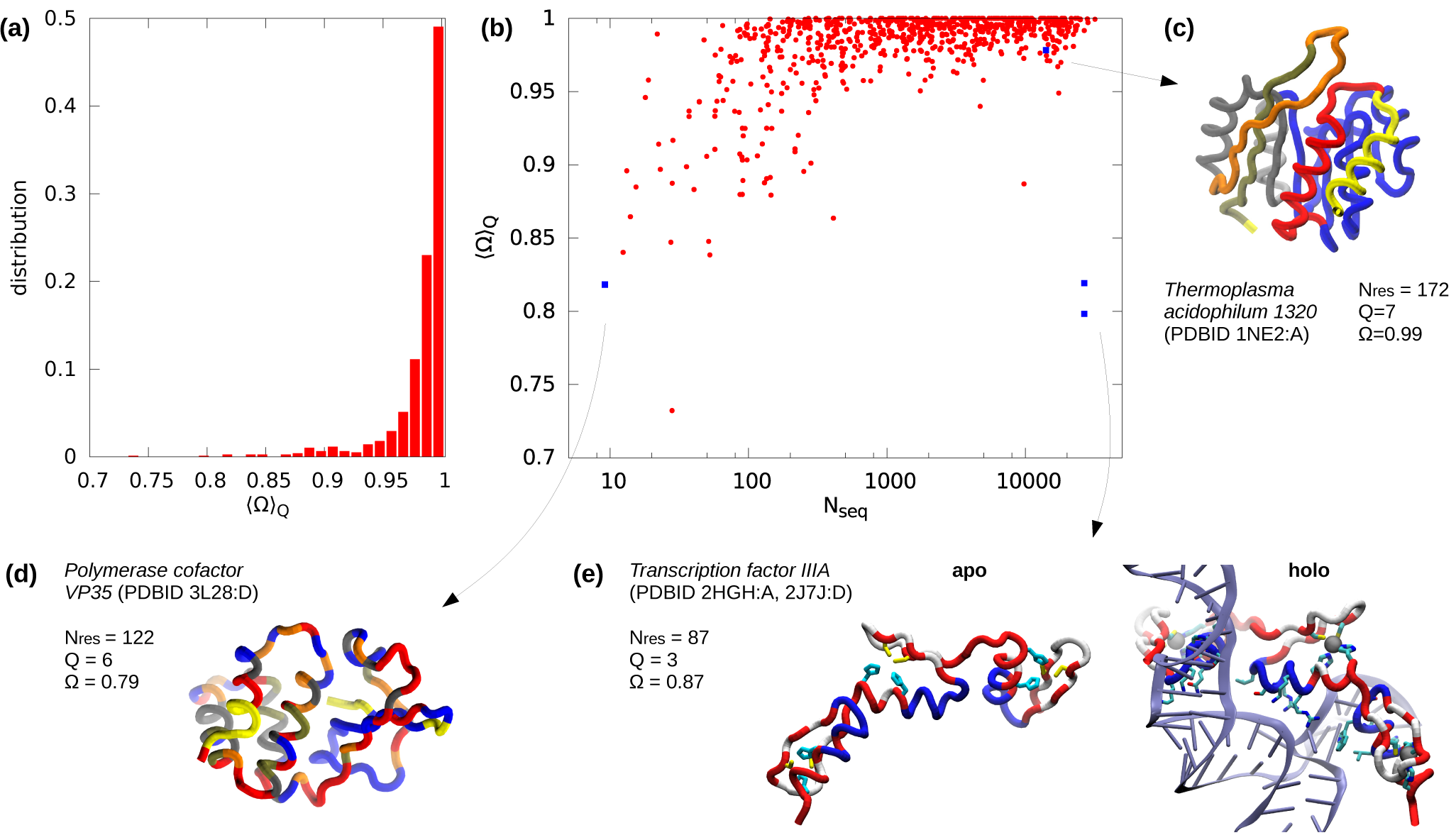
Histogram of the average structural compactness 〈Ω〉*Q* over the MSA dataset. (b) Scatter plot of the EDstructural compactness Ω for each single MSA α, averaged over the subdivisions into *Q =*2,…, 10 domains, *vs* the MSA size. (c-e) Structural representation of three notable examples of ED decompositions highlighted in (b). (e) Views of the Transcription factor IIIA (PDBID: 2HGH), in the apo form and in complex with a 5S rRNA 55mer

The average data in Fig. 2, as well as in Fig. S5 and S6, indicate that throughout a wide range of subdivisions, the EDs for the various MSAs retain a very good degree of compactness. The distribution in Fig. 2a is, in fact, clearly skewed toward 1, and the median value is as large as 0.99. It is also noteworthy that, for a given protein, the degree of spatial compactness is almost constant upon increasing the number of subdivisions, see Fig. S7. This means that the method identifies structurally meaningful subdivision for several levels of coarse graining, from low to high resolution.

As for the clustering coefficient, the plot in Fig. 2b highlights the correlation between the compactness and the number of sequences in the MSA. It is seen that the lowest (though still high) degree of compactness are typically found for MSAs with only 300 sequences or less. We interpret this result as an indirect indication that, when less than 300 sequences are used to infer the couplings, the network is not reliably reconstructed and consequently the ED subdivisions are less compact.

To illustrate the concepts discussed above within the context of selected protein structures, we show in Fig. 2c-e few examples of ED subdivisions.

The example in Fig. 2c corresponds to a MSA with a large pool of sequences (14,080) and an average compactness 〈Ω〉_*Q*_ = 0.98. The structure shown in the figure is the representative PDB entry 1NE2 and the subdivision corresponds to the optimal partitioning (*Q* = 7). Its high degree of compactness, Ω= 0.99, is readily perceived by inspecting the subdivisions that, with the sole exception of a terminal residue, are visibly compact along the primary sequence and in space too. The other two examples in Fig. 2d-e pertain to two distinct instances whose average compactness is about 0.8, *i.e.* on the low side of the distribution. The first instance is for the Ebola viral protein 35, represented by PDB entry 3L28, which has the least numerous MSA in the data set, involving 9 sequences only. For this limiting case with very poor statistics, the shown optimal subdivision presents a noticeable fragmentation for each of the *Q* = 6 domains. The other case is arguably a more interesting outlier, because it corresponds to one of the most numerous MSAs. Specifically, it corresponds to the transcription factor IIIA (TFIIIA), a Cys2His2 zinc finger protein involved in nucleic acid recognition and regulation [63]. The two structures in Fig. 2e pertain to the fingers 4-6, in the free state [64] (2J7J), and bound to 5S rRNA 55mer [65] (2HGH), the latter having two more residues in the C-terminus. The optimal partitioning involves *Q* = 3 EDs. When such subdivision is applied to the apo structure of the zinc finger, it appears fragmented due to the mutual interspersing of stretches assigned to different domains. In contrast to the previous case in Fig. 2d, however, the residues belonging to the three domains are not randomly distributed and show a well-defined structural organization. Specifically, the partitioning of a single zinc finger is repeated identically for all the three motifs. Indeed, when the same subdivision is applied to the RNA bound form in Fig. 2e, one sees that the domains acquire a more meaningful spatial organization. In fact: (i) the red domain defines the binding site formed by two cysteines on the *β*-hairpin and two histidines in the helix (highlighted in the apo form in yellow and cyan, respectively) that coordinate the zinc ions crucial to stabilize the fold [65]; (ii) the white domain sustains and locks the hairpin onto the helix (note the facing white residues, consistently present in all three helices); and, finally, (iii) the remaining blue part of the helix (referred to as “recognition helix”) contains residues forming sequence-specific contacts with the groove of the nucleic acid.

It is therefore remarkable, and intriguing, that the fragmented nature of this outlier is accidental, since the divisions acquire a clearer—and functionally-oriented—meaning when viewed in the context of the holo form. This further underscores the functional significance of EDs based on the MSA analysis.

### Correlation with Dynamical Domains

Motivated by these observations, we undertook a systematic comparison of the EDs and the quasi-rigid (or dynamical) domains (DDs) for each of the 813 MSAs. Specifically, the DDs were obtained by an elastic network (ENM) analysis [66, 67] of the reference structure for each MSA [32]. The structure-based character of the DD analysis is an apt complement of the sequence-based one of EDs. This duality makes the comparison particularly interesting and relevant for framing the sequence → structure → function relationships.

The overlap of the two types of domain subdivisions was measured in terms of the adjusted mutual information (AMI), which allows for a straightforward assessment of the statistical significance of the subdivisions overlap, see Methods.

To better illustrate the correspondence of the EDs and DDs, which we recall are established respectively using sequence-and structure-based features, and to give an immediate meaning to the AMI value, we discuss here two examples, the SbmC protein and the ABC transporter. Fig. 3a shows the SbmC protein (PDB code 1JYH, 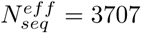) for which the *Q* = 4 ED decomposition overlaps almost perfectly with the dynamical one (AMI=0.8), grouping together not only secondary structure elements but also tertiary contacts. It is interesting to note that, as in Fig. 1b, the two clusterings share similar patterns for the quality score, and that all the subdivisions display a relatively large AMI. Also for the example in Fig. 3b (ABC transporter, PDB code 2ONK, 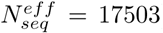), a consistent overlap between EDs and DDs is observed at all levels of protein subdivision. In particular we note that also the lowest AMI value of 0.5 (for *Q* = 4) shows already a highly satisfactory agreement between the two partitionings. In Fig. 3c we extend the analysis to the entire database, analyzing, for each dataset, both the largest AMI (*AMI_max_*) and the average one (〈*AMI*〉*Q*) over the interval *Q*=2,…, 10. We observe again a strong dependence on 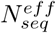: for MSAs with 500 sequences or more, the average values for *AMI_max_*and 〈*AMI*〉*Q* are 0.62 and 0.47, compared to the corresponding values of 0.49 and 0.35 when 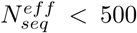, respectively.

**Figure 3.**
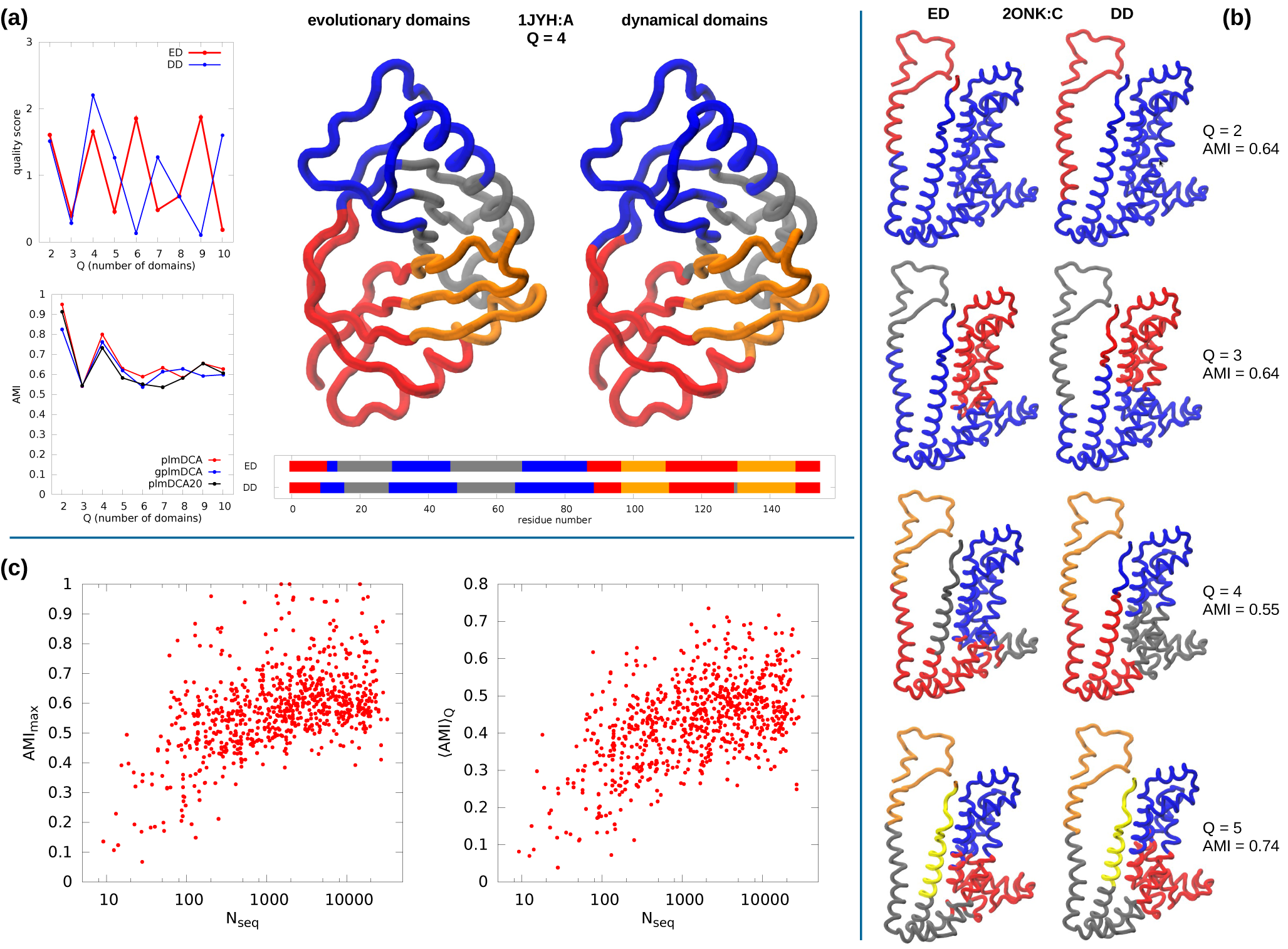
(a-b) Evolutionary (ED) and dynamical (DD) domain decompositions of a SbmC protein (PDB ID: 1JYH:A) and an ABC transporter permease protein (PDB ID: 2ONK:C). (c) Scatter plots of the maximum (left) and average (right) adjusted mutual information, over the domain number *Q,* with the dynamical decompositions, as a function of the effective MSA size.

The remarkable overlap between EDs and DDs at *all* levels of subdivision suggests that our clustering approach captures all the relevant topological features of the network of statistical dependencies and thus constitutes a potentially powerful tool to investigate, for instance, the sequence determinants of functional differentiation within a given protein architecture. In this comparative scenario, the small number of sequences available for each functional group does not allow a reliable quantitative estimation of the coevolutionary couplings; however, thanks to their robustness, EDs can be confidently defined and compared across distinct phylogenetic groups.

## Case Study: Comparative Analysis Across the 6TM Family of Ion Channels

We conclude by applying the ED analysis to a specific class of ion channels, the six-transmembrane-helices (6TM) superfamily, for which the sequence-function relationship has been actively investigated in a number of seminal studies [68].

For definiteness we focus on three different 6TM families: the voltage-gated potassium-selective channel (Kv, PDB code 2R9R [69]), the bacterial voltagegated sodium-selective channel (BacNav, 4EKW [70]) and transient receptor potential (TRP, 3J5P [71]) channels. These families are characterized by a strictly conserved tetrameric architecture. The latter is shown in Fig. 4a where different colors are used to highlight the main functional domains, including the four-helix bundle voltage sensor domain (VSD) and the pore of the ion conduction pathway, which involves two trans-membrane helices and the linking reentrant pore, containing the selectivity filter.

**Figure 4.**
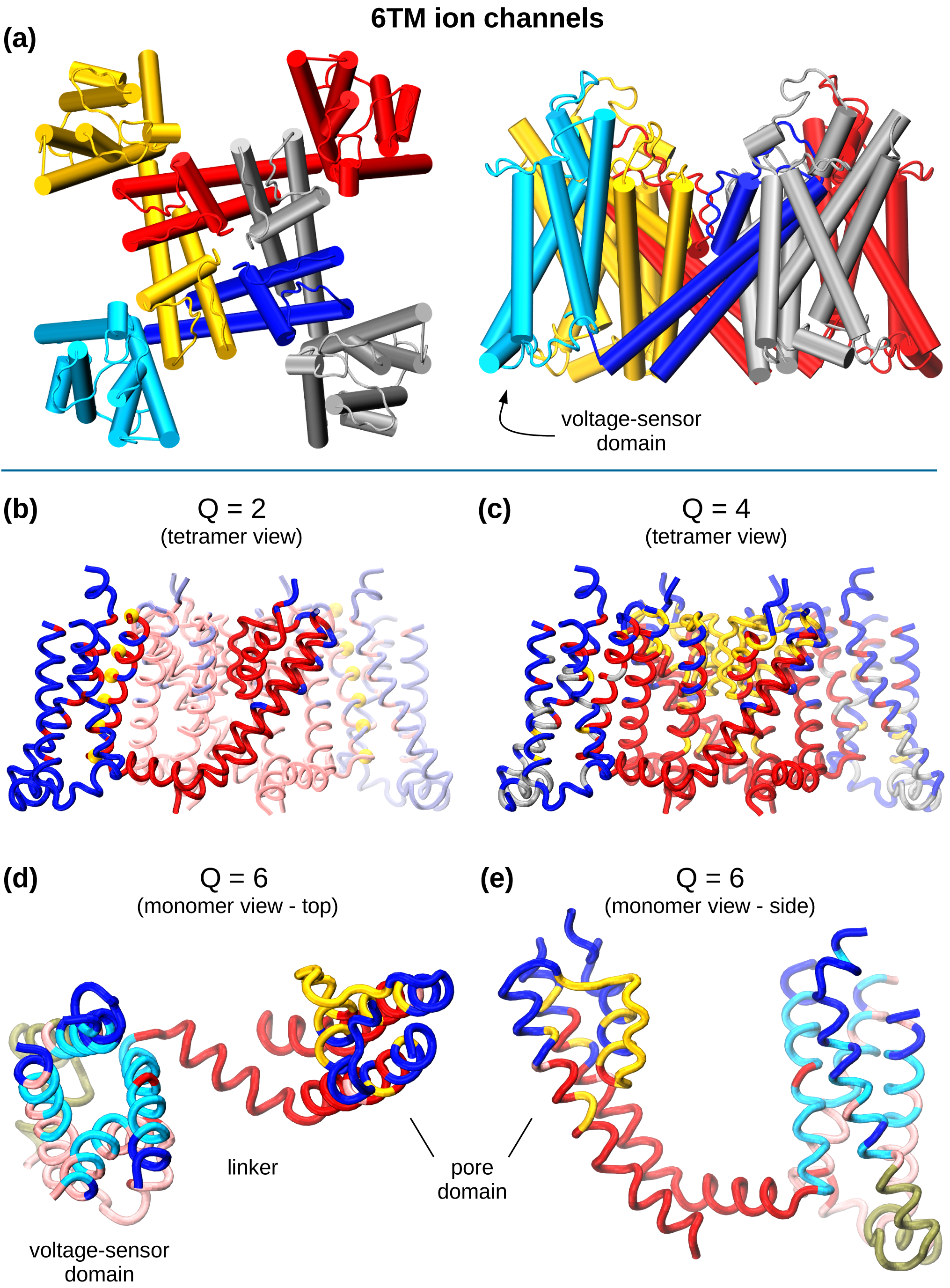
Evolutionary domains for Kv channels. a) Schematic representation of biological tetrameric assembly of 6TM-channels, with each color representing a single monomeric subunit (top and lateral view). For the blue subunit, the voltage sensor domain is highlighted in cyan. b) Representation of the most meaningful subdivisions (see also Fig. S11 for the quality score) on the tetrameric assembly. For Q=2 the positive charged residue responsible for voltage sensing are reported as yellow spheres.

This single structural template inherited from an ancestor gene has enabled, through differentiation, an explosion of functional variability. Channels in the 6TM class, in fact are involved, for instance, in reporting noxious environmental conditions, in shaping the neuronal action potential and in syncing the beating of the heart [59]. Since all these channels share the same architecture, different decompositions in EDs in different phylogentic groups likely reflect distinct functional rather than structural aspects [51, 52].

We analyzed the MSAs for the three families based on a pool of ∼ 800 sequences, each with 200 positions [72] from which we omitted the highly gapped regions of the alignments (typically occurring in loops between the six trans-membrane helices). Although the dataset that we used is the most comprehensive available at the moment, its size is clearly limited by comparison to the much better populated cases discussed in the previous sections, showing a pretty low ∆*C* (see Fig. S10). To ensure a robust analysis, we decided to decompose the graph corresponding to the maximum ∆*C* for each MSA.

In Fig. 4b we report various subdivisions for the Kv family for increasing numbers of domains (see Fig. S11 for quality score). The subdivision for *Q* = 2 is already unexpectedly informative, since the the fourth helix of the VSD (called S4) and its facing residues, are associated with the pore domain rather than the rest of the VSD. This is an intriguing result because the aforementioned classic subdivision into structural domains would have kept these elements apart.

From a functional point of view, however, the sequence-based subdivision of the primary (*Q* = 2) EDs is meaningful. In fact, it agrees with the strong mechanical coupling between the pore region and S4 [73–75]. We recall that the latter contains the positive residues (yellow spheres in Fig. 4b) that sense trans-membrane potential variations and determine the movement of this helix across the membrane; this movement is, in turn, transmitted to the pore domain for gating.

The division into *Q* = 4 EDs, in Fig. 4c, picks up further functional features. One domain largely corresponds to the selectivity region, formed by all the residues lining the narrow and highly conductive ion pathway (in yellow), another is associated to the gating region (red) and the other two comprise respectively the internal and external residues of the VSD. It is notable that a different domain assignment is found for the two faces of the pore helix, with the upper one sustaining the selectivity filter and the lower one contacting the gating domain.

As anticipated in the previous sections, this Kv example is also particularly instructive of the multi-level description that EDs can provide about the various protein features. Finer subdivisions (*Q* = 6) mostly return the basic structural elements of the system. In this subdivision the pore and the voltage sensor regions are mostly assigned to different EDs, with the exception of the extracellular portion of the channel (highlighted in blue in Fig. 4d-e) which still bridges between the two. When viewed in the context of the channel tetramer, it appears natural to speculate that this region is instrumental for the signal propagation between the loops of the voltage-sensing and the pore domains, which can indeed be modulated by external stimuli, like ligand binding. We accordingly surmise that amino acids in this region are genuinely evolutionary-related for functional reasons.

Further elements regarding the functional role of EDs emerge from the comparison of Kv, BacNav and TRP subdivisions, which are given in Fig.5 and are further detailed in Fig. S11.

**Figure 5.**
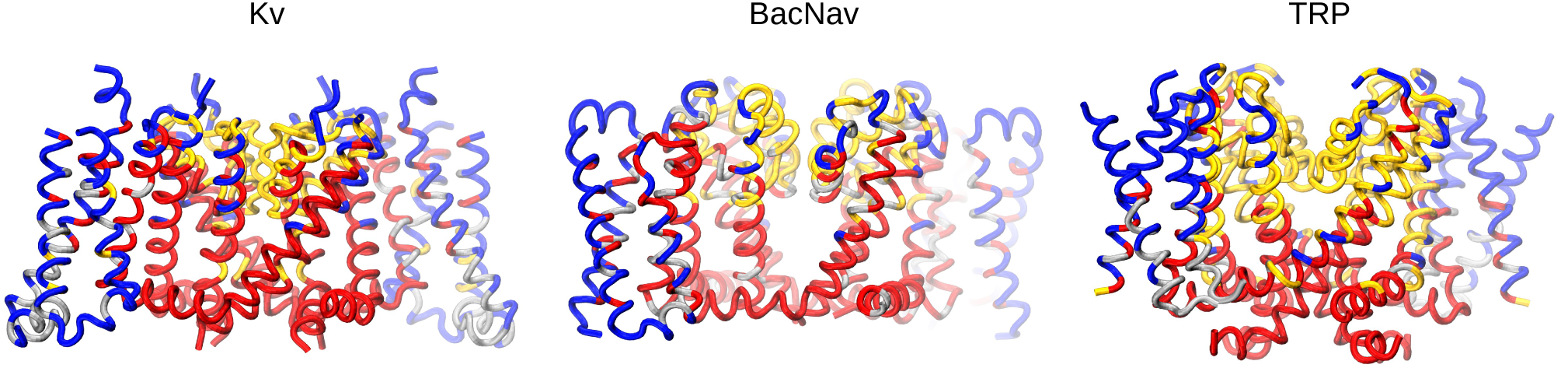
Comparative analysis of the Evolutionary Domains for KV, BachNav and TRP channels, corresponding to subdivision Q=4 (see quality score in Fig. S11). While Kv and BacNav show similar organization, coherent with their analogous functional requirements, TRP is characterized by a different domain pattern, consistently with its ligand-gated properties and loss of voltage-gated ones, specific of the other two channels.

The comparison between Kv and BacNav (another tetrameric voltage-gated family, selective for sodium) reflects how the functional constraints shaped these two families along evolution, in an almost superimposable way. In fact, the S4 helix segregates with the lower part of the pore and together form the “gating domain” (in red). Similarly the reentrant pore helix is splitted into the upper and lower faces, sustaining the selectivity domain (in yellow), and the rest of VSD is grouped into internal and external residues.

The organization of EDs for TRP channels is instead totally different. Indeed this channel family, identified only in Eukaryota, has distinct characteristics with respect the other ones. Specifically, it is a non-selective cation channel gated by a variety of stimuli, such as temperature, pH and ligands binding [76–79]. In particular these channels have been shown to possess two different gating regions [71, 80], which are indeed well-captured by the ED decomposition. The division of S4 is compelling in this respect since it is consistent with the lack of the dynamical role that, instead, characterizes it in voltagegated ion channels: only the C-terminal residues are associated with the gating domain (in red). The upper part of S4 is instead longitudinally sectioned, with the internal residues all grouped with the upper part of the rest of the VSD. The external part of S4 belongs to the extended yellow domain: the latter represents effectively a second upper gating domain, as suggested in [71, 80]. Remarkably, the yellow cavity determined by the two pore helices and the external part of S4 correspond exactly to the location of the vannilloid pocket [80–84], which represents the main intracellular binding site for the activators of these channels.

## Conclusions

Patterns of correlated mutations in a multiple sequence alignment can be used to reveal a set of pairwise statistical interactions that are often informative about the possible spatial proximity between the residues involved. Strikingly, we showed that this network of couplings has a peculiar structure with large communities of residues that are more connected among themselves than with the rest of the sequence. Thus, beyond compensatory mutations involving pairs of contacting residues, entire groups of proteins residues appear to evolve in a concerted fashion. We characterized these communities, that we term evolutionary domains (EDs), by interpreting the statistical couplings as a measure of evolutionary proximity between residues. To this end, we used an efficient clustering framework, namely spectral clustering. When analyzed in the context of the protein structure, these couplings show an innate tendency to segregate into spatially localized and compact groups. We explored the possible biological meaning of these subdivisions, contrasting EDs with the dynamical, quasi-rigid domains identified by a recently introduced approach [32]. We found a large overlap between the two decompositions. Remarkably, the fact that the two clustering approaches provide consistent results for both small and large number of clusters is a strong indication that the method is accurate for a wide range of “spatial resolutions”: it highlights global functionally-oriented features when few large clusters are considered and it characterizes in details the local structure when a large number of clusters are analyzed. Therefore, even when the structure is unknown, our approach provides valuable information concerning both structure and function. In these cases, detecting the hierarchical organization in domains can represent a crucial initial step for any structural modeling with an atomistic level of detail. Even more interesting is the perspective of using this approach to engineer existing proteins: transferring EDs across proteins sharing the same architecture might enable *de novo* design of protein chimeras with novel biological properties.

Finally, despite the fact that DCA analysis depends crucially on the effective number of sequences used for the statistical inference, we observe robust and consistent results even for few hundreds of sequences. Thus EDs, and therefore the topology of the network of coevolutionary couplings, are more robust with respect to the sample size than coevolutionary coupling strengths [42, 43]. Importantly, this aspect widens *de facto* the scope of applicability of these inference methods beyond those cases for which thousands of sequences are available. In particular, this enables comparative studies in which homologous subfamilies, rather than the entire group, are studied separately with the ultimate goal of highlighting the functionally features distinctive of each subgroup, as shown here for the challenging case of ion channels.

## Material and Methods

The analysis described in this paper, and illustrated in Fig. 1a, is based on the interpretation of statistical couplings *J_ij_*, obtained by direct coupling analysis (DCA) of a protein MSA, as a measure of evolutionary proximity between residues *i* and *j*. Such similarity measure is then processed by a spectral clustering algorithm, which returns the most meaningful partitioning of the sequence into an optimal number of domains. The high robustness of the results has been investigated contrasting the subdivisions between three different DCA approaches and with respect to the dataset size (number of sequences).

## Direct coupling analysis

This term indicates a family of methods to predict contacts between amino acid pairs from the analysis of correlated mutations between sequence positions in a MSA.

The main characteristic of this analysis is the idea of disentangling direct from indirect couplings between sequence positions. By using a global statistical approach, in fact, it is possible to recognize the cases in which a high correlation between two residues is due to the fact that both of them depends statistically from a third variable.

A common starting point of these methods [40, 85] consists in considering each sequence *s_i_* of a MSA as one of the possible realizations from a probability distribution described by a Potts model:

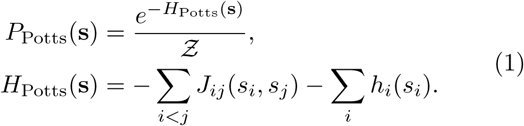

Each entry of the sequence *s_i_* can take on one of 21 symbols, corresponding to the 20 amino acid types (A, R, N, …), plus the gap symbol (-). The unknown parameters of the model, namely the couplings *h_i_*(*s_i_*)∈ℝ^21^and *J_ij_*(*s_i_,sj*) ∈ ℝ^21×21^, are then inferred based on a principle of maximum likelihood. The Frobenius norm of couplings || *J*_*ij*_(*s_i_,sj*)||, is finally taken as an estimate of the strength of the correlation between the two positions *i* and *j* on the sequence.

The problem of inverting this Potts model has been addressed in many ways [40–42]. In this paper, we referred to the pseudo-likelihood minimization approach (plmDCA) described in [86]. In recent years, this method has been further improved by including corrections aimed to cure the problem of mispredictions in presence of long stretches of gaps in the MSA sequences. In our discussion we considered two modifications to the original plmDCA scheme, introduced in [43], namely gplmDCA (gap-enhanced plmDCA) and plmDCA20. The former approach takes into account the effect of gap-rich sequence positions by including additional parameters in the Potts model; the latter simply excludes the contribution of the gap term when computing the Frobenius norm.

## The dataset

For our analysis we used the extensive dataset of 813 MSAs compiled in [43]. These alignments were computed by using the homology detection method HH-blits [87] starting from the sequence of a target PDB structure, and are characterized by a heterogeneous number of sequences (16–65535) and positions (30–494). The database was specifically aimed at testing the contact prediction capability of DCA methods. The relative structures have been used here *a posteriori* to examine the structural and dynamical characteristics of EDs that we infer on the basis of the MSA information only. For constructing the MSA for adenylate kinase we use the same approach described in [43]. Finally, the three ion-channel MSAs were the same produced and analyzed in [72].

## Spectral clustering

The clustering strategy adopted in this context is the spectral clustering algorithm, described for instance in [88]. This algorithm takes in input a matrix of pairwise similarities *S_ij_* between the elements to be clustered and returns a set of subdivisions into a variable number of clusters *Q* = *Q*_min_,…,*Q*_max_. This is done by performing a nonlinear mapping of the original data into a space of reduced dimensionality, where the strongest local similarity relationships between the elements are enhanced over the weakest ones, regardless of the global arrangement of the elements. Such dimensionality-reduction approach closely resembles the operation of finding the metastable states of a random walk, defined by a transition probabilities matrix [60, 89]. As it is often the case in clustering schemes, the optimal number of subdivisions, or domains, is generally not known *a priori.* Therefore first we identify the optimal subdivisions for a pre-assigned number of domains, *Q*, varying between 2 and 20. Next we introduce a quality score, whose maxima highlight the most meaningful subdivisions, according to the observed eigenvalue gaps in the spectrum of the Laplacian matrix used by the spectral clustering [88]. This clustering algorithm has been successfully applied, in a previous work by some of us [32], to the identification of groups of residues sharing common dynamical traits. For the purpose of this work, the similarities between the residues are derived from the couplings strength: the output values of the DCA methods have been shifted by the minimal value in order to guarantee that the similarities were all positive, and then squared in order to increase the separation between weak and strong similarities, as commonly done in spectral clustering.

## *k*-nearest neighbor graph and clustering coefficient

In the framework of spectral clustering, the pairwise similarities *S_ij_* are used for building a graph representation which maps the initial similarities into local neighborhood relationships between the graph vertices [60]. This is usually done by either defining a cutoff value for the similarities or choosing a maximum number of edges connecting each vertex to its nearest-neighbors. For example, for structure-based domain decompositions, like the one employed in section and in the analysis of adenylate kinase’s test case, the set of neighbors is naturally defined on the basis of the contact map of the protein, according to the strategy described in a previous work by some of us [32]. Here, since the clustering relies solely on sequence information, we do not use this structural information and adopt instead a symmetric *k*-nearest neighbor graph: for each residue only the *k* strongest couplings are kept. In order to always work on a completely connected graph, we also force the inclusion of edges between consecutive residues on the sequence.

To choose *k*, we use an internal criterion based on the clustering coefficient *C*[90], also known as “cliqueness”. This quantity measures the average probability that two neighbors of a vertex are also connected with each other. More precisely, given a vertex *v_i_* with *n_i_* neighbours, the local clustering coefficient is computed as 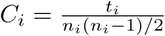, where *t_i_* is the number of links between the neighbors of *v_i_*. The global clustering coefficient *C* is then defined as the average of the local coefficients of vertices with more than one neighbor.

For comparison, it is informative to derive the clustering coefficient for random graphs, also known as Erdös-Rényi graphs. In these models, the probability *p* of edge formation is defined *a priori,* and it is independent for each vertex, so that the clustering coefficient is simply equal to *p*[91]. Given the number *N* of vertices, the average number of neighbours (or average degree) *k̅* is easily computed as the total number of edges, *N*(*N*− 1)/*p*, divided by *N*,*i.e. k̄*= (*N*− 1)*p*. It then follows that *C*_rand_=*p* = *k̄*/(*N* − 1). The difference of the clustering coefficient with respect to the random case, ∆*C*=*C* −*C*_rand_, provides then a good measure for the intrinsic clustering “propensity” of a graph. In the next section, we explain how this criterion can be adopted to guide the choice of the parameter *k*.

## Clustering Propensity of Couplings

In order to characterize the intrinsic clustering propensity of coevolutionary couplings and to investigate the role of the parameter *k*, for each entry in the database of 813 MSAs used in the Results, we produced the graphs *G_k_* for a wide range of *k* (3, 5, 7, 10, 15, 20, 25, 30, 40, 55). We then evaluated the optimal *k* by calculating the weighted clustering coefficient of each graph [62]. The histograms in Fig. S2 shows the *k* that maximizes the clustering coefficient ∆*C*=*C*–*C_rand_* for each MSA, for the three DCA methods. In most of the cases the optimal value is *k* = 7, for all the three coevolutionary analysis methods. This is true also when ∆*C* is averaged over the different MSAs and is analyzed as function of *k* (Fig. S3a). We notice that also the values *k* = 5 and *k* = 10 produce a relatively large value of the clustering coefficient, which, however, decreases fastly when *k* is greater than 10. Importantly, the distribution of the optimal *k* becomes increasingly more peaked around *k* = 7 (Fig. S2 in SI) with larger values of the clustering coefficient as the effective number of sequences increases (this number, denoted as 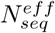, is the number of sequences whose maximum mutual identity is 90%), as clarified also by their strong correlation in fig. S4 for all DCA methods. The fact that *k* shows a definite value and that this value potentially results from a physical constraint (average number of structural neighbors) suggests that a parameter-free algorithm (with a fixed value of *k* = 7) is able to reliably detect the cluster structure of the most generic MSA. Accordingly, in the Results section, we always set *k* = 7.

## Structural compactness

The spatial compactness of the EDs on the protein structure has been assessed by introducing a new quantity, computed on the graph defined by the contact map between *C*_α_’s distant less than 10 Å. This quantity, that we call structural compactness (Ω), relies on the calculation of graph distances between residues belonging to the same domain. In case of a compact domain *q* of *n_q_* residues, each residue can be connected with any other one in the same domain by at least one path hopping only between members of the same cluster. When a single residue i is completely disconnected from the relative domain, the number of intra-cluster “broken paths” *b*_*q*_ is equal to *n_q_*−1 (counting symmetric paths once). We then say that there are *d_q_* = *b_q_*/(*n_q_* − 1) disconnected residues, which in the latter case amount to 1. We note that this definition distinguishes between a “fuzzy” domain, where the disconnected residues are scattered and do not group together, and a domain split in multiple sub-domains. In particular, the case in which half of the cluster members are scattered over the graph in isolated positions has a worse count (*b_q_*= (*n_q_*/2)(*n_q_* − 1), *d_q_*=*n_q_*/2) than the case where a cluster is split in two compact sub-clusters (*b_q_*= (*n_q_*/2)^2^,*d_q_*≃*n_q_*/4). The compactness Ω of a specific subdivision in *Q* is finally defined as

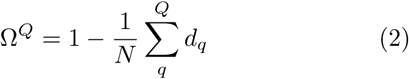

where *N* is the total number of residues.

## Adjusted Mutual Information

In clustering, when comparing two different partitionings of the same system, a possible way for quantifying their degree of similarity is to compute their mutual information (MI). In the context of information theory, the MI measures the amount of information shared by the two partitionings. Throughout our analysis, we make use of the related concept of adjusted mutual information (AMI) [92]. This quantity is given by 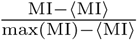, where 〈MI〉 is the expected value of MI over pairs of random partitions, for which an analytical expression has been derived [93]. The AMI has two desirable properties: (i) it is normalized, *i. e.* two identical partitions return an AMI equal to 1, and (ii) it is adjusted-for-chance, namely two random clusterings produce on average an AMI equal to 0. For normalization we take max(MI) as the minimal Shannon entropy between the two sets. This measure is conceptually similar to the Adjusted Rand Index, but it is usually more reliable in comparing subdivisions with different and high number of clusters [92].

## Robustness With Respect to the Dataset Size

One of the main limitations of coevolutionary methods is that their performances deteriorates sensibly with decreasing the number of available sequences [42, 43], undermining the accurate detection of statistical couplings and, consequently, of contacts. Also in our case the number of sequences plays a role in determining the clustering propensity of the similarity graph *G* (see fig. S4), but such an effect appears to be more gentle on the definition of EDs, as shown by the compactness analysis (Fig. 2b, S5 and S6). Here we investigate systematically the dependence of the clustering accuracy with the number of sequences by comparing the subdivisions of an MSA with the ones produced by two reduced MSAs (MSA’ and MSA”), containing, respectively, 10% and 1% of the sequences present in the original dataset. We repeated this calculation for four MSAs (PDB codes: 1NE2, 1XDZ, 2ONK, 3M0Z) characterized by the largest number of sequences in the database (*N*= 65535), obtaining the following average effective numbers of sequences: 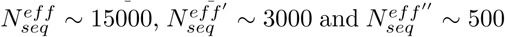. To quantify the similarity between two different partitionings, we use the AMI, described above. We summarize the analysis on these 12 datasets in Fig. S8 and S9, measuring, in particular, the AMI between the original and the reduced MSAs at varying number of clusters *Q*, together with the relative compactness and behavior of the quality score. Fig. S8 confirms how the prediction precision (PPV, measured using the same criteria as in [43], from the ratio of true contacts within the first strongest couplings considered) highly deteriorates for the reduced datasets, with the most dramatic effect for the poorest ones. Conversely, the EDs in all the cases show a remarkably high correlation at all levels of subdivision *Q* and for all dataset sizes. To get an immediate idea of the expected overlap for each AMI value, compare how consistent the decompositions characterized by an AMI value of 0.55 are in Fig. 3b; the best case scenario (bottom row, with an AMI value of 0.75) shows a visually indistinguishable degree of overlap. Also the compactness stays very high in all the datasets. We also note that the cases which show a lower AMI or compactness, especially at small *Q*s, are associated with the lowest values of the quality score. Moreover, peaks in the quality score indicate the subdivisions with the best overlap and compactness. Finally, the fact that the three datasets (and full and the reduced ones) show very similar profiles of the quality score, which highlight often the same subdivisions as most meaningful, shows, once again, how the three coupling matrices carry consistent information even when the statistics is extremely poor.

## Comparison and robustness for different inference methods

Each of the analyses contained in Results, benchmarking the performance of our coevolutionary clustering method, has been conducted for all three inference techiniques considered, and reported for plmDCA20 and gplmDCA in Supporting Information (SI). In Fig. S12 we produced the equivalent plot as in Fig. 3c. Also these two methods realize a high agreement between the derived EDs and the dynamical domains both for *AMI_max_* and 〈*AMI*〉*_Q_*, totally correlated with the number of sequences available for the inference. Figs. S13-S15 show that the performances of the three methods in terms of overlap with the dynamical domains (DDs) appears even more similar than compared with the clustering coefficient and compactness, outlining the stability and reproducibility of EDs with respect to the initial definition of the couplings. The strict consistency between the divisions is also confirmed from their strong correlation in the *AMI_max_* and 〈*AMI*〉_Q_ values with the DDs (see Fig. S14) and from the high reciprocal AMI values (Fig. S15) when we compare directly the EDs obtained by the three methods, obviously correlating with 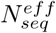. Based on the correlation with the DDs we have also checked *a posteriori,* in Fig. S16, that the choice of *k*= 7 in place of a different *k_max_* doesn’t affect much the results: in general the average AMI increases decisively when *k_max_* was larger than 15, and slightly decreases when *k_max_* was smaller than 7, especially in association with case with low 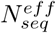, when fewer strong couplings reflects true contacts. As a final comment on the comparison, in all the analyses the gplmDCA approach appears to perform worse than the other two, showing in particular many datasets at high 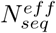 having lower clustering coefficient and lower AMI values both with DDs and with the EDs from the other methods. The differences instead between plmDCA and plmDCA20 appears really modest, especially taking in account the small advantage in compactness of the first (Fig. S5), which intuitively would favor a better correlation with the structurally-compact DDs. This is in a sense expected: the two methods do not differ in the inference technique (as for gplmDCA), but only in the derivation of the final scoring for the coupling *J_ij_*, since in plmDCA20 the contribution from gap variable is ignored. Nevertheless, as we discussed earlier and showed in Fig. S3, this small deviation can determine, in comparison, stronger couplings between adjacent residues for plmDCA. This usually leads basically to the same subdivisions, but in few cases we noticed differences in the EDs obtained from the two approaches: plmDCA-derived subdivisions seems to follow more the primary and secondary structure of the protein, while the plmDCA20-derived ones seems to outline better tertiary contacts between these elements. We underline that both perspectives can be equally important: the former can emphasize more the presence of different protein domains or modules originated, for example, from the gene fusion of two distinct proteins; on the other hand, the latter can inform more about the conserved tertiary assembly of such domains, possibly highlighting long-range contacts crucial for the protein function.

## References

[1] P. K. Agarwal, S. R. Billeter, P. R. Rajagopalan, S. J. Benkovic, and S. Hammes-Schiffer, Proceedings of the National Academy of Sciences 99, 2794 (2002).

[2] S. Hammes-Schiffer and S. J. Benkovic, Annu. Rev. Biochem. 75, 519 (2006).

[3] V. Carnevale, S. Raugei, C. Micheletti, and P. Carloni, Journal of the American Chemical Society 128, 9766 (2006).

[4] C. Chennubhotla and I. Bahar, PLoS Comput Biol 3, e172 (2007).

[5] V. Carnevale, F. Pontiggia, and C. Micheletti, Journal of Physics: Condensed Matter 19, 285206 (2007).

[6] A. Zen, V. Carnevale, A. M. Lesk, and C. Micheletti, Protein Science 17, 918 (2008).

[7] A. del Sol, C.-J. Tsai, B. Ma, and R. Nussinov, Structure 17, 1042 (2009).

[8] C. J. Jackson, J.-L. Foo, N. Tokuriki, L. Afriat, P. D. Carr, H.-K. Kim, G. Schenk, D. S. Tawfik, and D. Ollis, Proceedings of the National Academy of Sciences 106, 21631 (2009).

[9] K. Teilum, J. G. Olsen, and B. B. Kragelund, Cellular and Molecular Life Sciences 66, 2231 (2009).

[10] G. Morra, G. Verkhivker, and G. Colombo, PLoS Comput Biol 5, e1000323 (2009).

[11] D. Provasi, M. C. Artacho, A. Negri, J. C. Mobarec, and M. Filizola, PLoS Comput Biol 7, e1002193 (2011).

[12] G. Bhabha, J. Lee, D. C. Ekiert, J. Gam, I. A. Wilson, H. J. Dyson, S. J. Benkovic, and P. E. Wright, Science 332, 234 (2011).

[13] V. N. Bavro, R. De Zorzi, M. R. Schmidt, J. R. Muniz, L. Zubcevic, M. S. Sansom, C. Vénien-Bryan, and S. J. Tucker, Nature structural & molecular biology 19, 158 (2012).

[14] T. J. Glembo, D. W. Farrell, Z. N. Gerek, M. Thorpe, and S. B. Ozkan, PLoS Comput Biol 8, e1002428 (2012).

[15] Y. Liu and I. Bahar, Molecular biology and evolution 29, 2253 (2012).

[16] J. Lai, J. Jin, J. Kubelka, and D. A. Liberles, Journal of Molecular Biology 422, 442 (2012).

[17] C. Micheletti, Physics of Life Reviews 10, 1 (2013).

[18] E. Z. Eisenmesser, D. A. Bosco, M. Akke, and D. Kern, Science 295, 1520 (2002).

[19] E. Z. Eisenmesser, O. Millet, W. Labeikovsky, D. M. Korzhnev, M. Wolf-Watz, D. A. Bosco, J. J. Skalicky, L. E. Kay, and D. Kern, Nature 438, 117 (2005).

[20] W. Min, X. S. Xie, and B. Bagchi, The journal of physical chemistry B 112, 454 (2008).

[21] W. Min, X. S. Xie, and B. Bagchi, The Journal of chemical physics 131, 08B606 (2009).

[22] Z. Nevin Gerek, S. Kumar, and S. Banu Ozkan, Evolutionary applications 6, 423 (2013).

[23] D. E. Shaw, M. M. Deneroff, R. O. Dror, J. S. Kuskin, R. H. Larson, J. K. Salmon, C. Young, B. Batson, K. J. Bowers, J. C. Chao, et al., Communications of the ACM 51, 91 (2008).

[24] D. E. Shaw, P. Maragakis, K. Lindorff-Larsen, S. Piana, R. O. Dror, M. P. Eastwood, J. A. Bank, J. M. Jumper, J. K. Salmon, Y. Shan, and W. Wriggers, Science 330, 341 (2010).

[25] D. Granata, C. Camilloni, M. Vendruscolo, and A. Laio, Proceedings of the National Academy of Sciences 110, 6817 (2013).

[26] W. Zheng, B. R. Brooks, and D. Thirumalai, Biophysical journal 93, 2289 (2007).

[27] A. Zen, C. Micheletti, O. Keskin, and R. Nussinov, BMC structural biology 10, 26 (2010).

[28] A. Ramanathan and P. K. Agarwal, PLoS Biol 9, e1001193 (2011).

[29] S. Hayward, A. Kitao, and H. J. Berendsen, Proteins: Structure Function, and Genetics 27, 425 (1997).

[30] R. Potestio, F. Pontiggia, and C. Micheletti, Biophysical journal 96, 4993 (2009).

[31] G. Polles, G. Indelicato, R. Potestio, P. Cermelli, R. Twarock, and C. Micheletti, PLoS Comput Biol 9, e1003331 (2013).

[32] L. Ponzoni, G. Polles, V. Carnevale, and C. Micheletti, Structure 23, 1516 (2015).

[33] E. I. Shakhnovich and A. M. Gutin, Proceedings of the National Academy of Sciences 90, 7195 (1993).

[34] N. V. Dokholyan and E. I. Shakhnovich, Journal of molecular biology 312, 289 (2001).

[35] A. Sali, L. Potterton, F. Yuan, H. van Vlijmen, and M. Karplus, Proteins: Structure Function, and Genetics 23, 318 (1995).

[36] C. A. Rohl, C. E. Strauss, K. M. Misura, and D. Baker, in Methods in Enzymology (Elsevier BV, 2004) pp. 66–93.

[37] A. Roy, A. Kucukural, and Y. Zhang, Nature Protocols 5, 725 (2010).

[38] S. W. Lockless, Science 286, 295 (1999).

[39] J. Dutheil, T. Pupko, A. Jean-Marie, and N. Galtier, Molecular biology and evolution 22, 1919 (2005).

[40] M. Weigt, R. A. White, H. Szurmant, J. A. Hoch, and T. Hwa, Proceedings of the National Academy of Sciences 106, 67 (2008).

[41] F. Morcos, A. Pagnani, B. Lunt, A. Bertolino, D. Marks, C. Sander, R. Zecchina, J. Onuchic, T. Hwa, and M. Weigt, Proc Natl Acad Sci USA 108, E1293 (2011).

[42] H. Kamisetty, S. Ovchinnikov, and D. Baker, Proceedings of the National Academy of Sciences 110, 15674 (2013).

[43] C. Feinauer, M. J. Skwark, A. Pagnani, and E. Aurell, PLoS Comput Biol 10, e1003847 (2014).

[44] M. J. Skwark, D. Raimondi, M. Michel, and A. Elofsson, PLoS Comput Biol 10, e1003889 (2014).

[45] S. Hayat, C. Sander, D. S. Marks, and A. Elofsson, Proceedings of the National Academy of Sciences 112, 5413 (2015).

[46] J. Dutheil, T. Pupko, A. Jean-Marie, and N. Galtier, Molecular biology and evolution 22, 1919 (2005).

[47] D. A. Liberles, S. A. Teichmann, I. Bahar, U. Bastolla, J. Bloom, E. Bornberg-Bauer, L. J. Colwell, A. De Koning, N. V. Dokholyan, J. Echave, et al., Protein Science 21, 769 (2012).

[48] G. Süel, S. Lockless, M. Wall, and R. Ranganathan, Nat Struct Biol 10, 59 (2003).

[49] N. Halabi, O. Rivoire, S. Leibler, and R. Ranganathan, Cell 138, 774 (2009).

[50] R. S. Dwyer, D. P. Ricci, L. J. Colwell, T. J. Silhavy, and N. S. Wingreen, Genetics 195, 443 (2013).

[51] E. Palovcak, L. Delemotte, M. L. Klein, and V. Carnevale, The Journal of General Physiology 143, 145 (2014).

[52] E. Palovcak, L. Delemotte, M. L. Klein, and V. Carnevale, The Journal of General Physiology 146, 37 (2015).

[53] L. Sutto, S. Marsili, A. Valencia, and F. L. Gervasio, Proceedings of the National Academy of Sciences 112, 13567 (2015).

[54] K. N. Woods and J. Pfeffer, Mol Biol Evol 33, 40 (2015).

[55] M. Figliuzzi, H. Jacquier, A. Schug, O. Tenaillon, and M. Weigt, Mol Biol Evol 33, 268 (2015).

[56] A. Haldane, W. F. Flynn, P. He, R. Vijayan, and R. M. Levy, Protein Science 25, 1378 (2016).

[57] L. Sutto, S. Marsili, A. Valencia, and F. L. Gervasio, Proceedings of the National Academy of Sciences 112, 13567 (2015).

[58] A. F. Poon, L. C. Swenson, W. W. Dong, W. Deng, S. L. K. Pond, Z. L. Brumme, J. I. Mullins, D. D. Richman, P. R. Harrigan, and S. D. Frost, Molecular biology and evolution 27, 819 (2010).

[59] F. H. Yu and W. A. Catterall, Science Signaling 2004,re15 (2004).

[60] U. von Luxburg, Statistics and Computing 17, 395 (2007).

[61] M. Punta, P. C. Coggill, R. Y. Eberhardt, J. Mistry, J. Tate, C. Boursnell, N. Pang, K. Forslund, G. Ceric, J. Clements, A. Heger, L. Holm, E. L. L. Sonnhammer, S. R. Eddy, A. Bateman, and R. D. Finn, Nucleic Acids Research 40, D290 (2011).

[62] J. Saramäki, M. Kivelä, J.-P. Onnela, K. Kaski, and J. Kertész, Physical Review E 75 (2007), 10.1103/physreve.75.027105.

[63] J. Hanas, D. Hazuda, D. Bogenhagen, F. Wu, and C. Wu, J Biol Chem 258, 14120 (1983).

[64] D. Lu and “A. Klug, Invariance of the zinc finger module:a comparison of the free structure with those in nucleic-acid complexes,” (2007).

[65] B. M. Lee, J. Xu, B. K. Clarkson, M. A. Martinez-Yamout, H. J. Dyson, D. A. Case, J. M. Gottesfeld, and P. E. Wright, Journal of Molecular Biology 357, 275 (2006).

[66] C. Micheletti, P. Carloni, and A. Maritan, Proteins:Structure Function, and Bioinformatics 55, 635 (2004).

[67] E. Fuglebakk, N. Reuter, and K. Hinsen, J.Chem. Theory Comput. 9, 5618 (2013).

[68] B. Hille et al., Ion channels of excitable membranes, Vol. 507 (Sinauer Sunderland, MA, 2001).

[69] S. B. Long, X. Tao, E. B. Campbell, and R. MacKinnon, Nature 450, 376 (2007).

[70] J. Payandeh, T. Scheuer, N. Zheng, and W. A. Catterall, Nature 475, 353 (2011).

[71] M. Liao, E. Cao, D. Julius, and Y. Cheng, Nature 504, 107 (2013).

[72] M. Kasimova, D. Granata, and V. Carnevale, Curr Top Membr 78, 261 (2016).

[73] Z. Lu, A. M. Klem, and Y. Ramu, The Journal of General Physiology 120, 663 (2002).

[74] A. Broomand, R. Männikkö, H. P. Larsson, and F. Elinder, The Journal of General Physiology 122, 741 (2003).

[75] S. Long, E. Campbell, and R. Mackinnon, Science 309, 903 (2005).

[76] T. Voets, K. Talavera, G. Owsianik, and B. Nilius, Nature Chemical Biology 1, 85 (2005).

[77] I. S. Ramsey, M. Delling, and D. E. Clapham, Annual Review of Physiology 68, 619 (2006).

[78] Q. Feng, in Current Topics in Membranes(Elsevier BV, 2014) pp. 19–50.

[79] V. Carnevale and T. Rohacs, Pharmaceuticals 9, 52 (2016).

[80] E. Cao, M. Liao, Y. Cheng, and D. Julius, Nature 504, 113 (2013).

[81] F. Yang, X. Xiao, W. Cheng, W. Yang, P. Yu, Z. Song, V. Yarov-Yarovoy, and J. Zheng, Nature Chemical Biology 11, 518 (2015).

[82] L. Darré and C. Domene, Mol. Pharmaceutics 12, 4454(2015).

[83] K. Elokely, P. Velisetty, L. Delemotte, E. Palovcak, M. L. Klein, T. Rohacs, and V. Carnevale, Proceedings of the National Academy of Sciences 113, E137 (2015).

[84] Y. Gao, E. Cao, D. Julius, and Y. Cheng, Nature 534,347 (2016).

[85] A. Lapedes, B. Giraud, L. Liu, and G. Stormo, Correlated mutations in protein sequences: Phylogenetic and structural effects, Tech. Rep. (1998).

[86] M. Ekeberg, C. Lövkvist, Y. Lan, M. Weigt, and E. Aurell, Physical Review E 87 (2013), 10.1103/physreve.87.012707.

[87] M. Remmert, A. Biegert, A. Hauser, and J. Söding, Nature Methods 9, 173 (2011).

[88] A. Y. Ng, M. I. Jordan, Y. Weiss, et al., Advances in neural information processing systems 2, 849 (2002).

[89] B. Nadler, S. Lafon, R. R. Coifman, and I. G. Kevrekidis, Applied and Computational Harmonic Analysis 21, 113(2006).

[90] D. Watts and S. Strogatz, Nature 393, 440 (1998).

[91] M. E. J. Newman, in From the Genome to the Internet (Wiley-Blackwell, 2004) pp. 35–68.

[92] N. X. Vinh, J. Epps, and J. Bailey, in Proceedings of the 26th Annual International Conference on Machine Learning - ICML ’09 (Association for Computing Machinery (ACM), 2009).

[93] N. X. Vinh, J. Epps, and J. Bailey, Journal of Machine Learning Research 11, 2837 (2010).

